# Developmental biomechanics and age polyethism in leaf-cutter ants

**DOI:** 10.1101/2023.02.13.528272

**Authors:** Frederik Püffel, Lara Meyer, Natalie Imirzian, Flavio Roces, Richard Johnston, David Labonte

## Abstract

Many social insects display age polyethism: young workers stay inside the nest, and only older workers forage. This behavioural transition is accompanied by genetic and physiological changes, but the mechanistic origin of it remains unclear. To investigate if the mechanical demands of foraging effectively prevent young workers from partaking, we studied the biomechanical development of the bite apparatus in *Atta vollenweideri* leaf-cutter ants. Fully-matured foragers generate peak *in-vivo* bite forces of around 100 mN, more than one order of magnitude in excess of those measured for freshly-eclosed callows of the same size. This change in bite force was accompanied by a sixfold increase in the volume of the mandible closer muscle, and by a substantial increase of the flexural rigidity of the head capsule, driven by a significant increase in both average thickness and indentation modulus of the head capsule cuticle. Consequently, callows lack the muscle force capacity required for leaf-cutting, and their head capsule is so compliant that large muscle forces may cause damaging deformations. On the basis of these results, we speculate that continued biomechanical development post eclosion may be a key factor underlying age polyethism, wherever foraging is associated with mechanical demands on the musculoskeletal system.

## Introduction

Social insects are extremely ‘successful’ [1, 2], and this success is thought to be partially based on the evolution of a ‘division of labour’; some tasks are preferentially or exclusively performed by specific individuals. In many social insects, such task preferences transcend the elementary dichotomy between reproductive and non-reproductive labour, and sterile workers show preferences and specialisation for subsets of non-reproductive colony tasks. The classic explanation for this phenomenon suggests that task specialisation increases the ergonomic efficiency of colonies, and thus their fitness [e. g. 3, but see 4]. Two main themes are common to studies which propose explanations for the benefits *of* or possible mechanisms *for* the evolution of a non-reproductive division of labour [e. g. 5–8]: task preferences are associated with differences in *worker phenotype*, for example in terms of worker size or body shape, or vary with *worker age*.

In social insects, systematic changes in task preferences with age, or age polyethism, have evolved in bees [9–13], wasps [14, 15], ants [16–25], and termites [26]: freshly-eclosed workers tend to stay inside the nest and attend to queen and brood, and only older workers engage in foraging tasks outside the nest. This behavioural transition typically occurs within the first few weeks after eclosion [e. g. 9, 22, 23, 26], although the exact timeline is subject to variation based on genotype [20] and colony size [21, 27].

Because of its frequent emergence and importance to the ecology of social insects, age polyethism has received considerable attention from behavioural biologists [13, 15, 20, 21, 27], ecologists [14, 28, 29], geneticists [30, 31], neuroethologists [22, 32, 33], and biomechanists alike [34, 35], and several genetic and physiological correlates have been identified. For example, the transition from within-nest to outside foraging tasks is accompanied by substantial changes in gene expression [30, 31, 36], hormone and neuropeptide levels [14, 15, 27, 37], the exocrine system [13], muscle chemistry [38–40], and brain physiology and size [33, 41].

In particular, the physiological development following eclosion is hypothesised to be key in determining the ability of workers to perform specific tasks [7, 12, 22, 24, 32], suggesting an emergence of age polyethism based on the acquisition of new capabilities [22, but see 6, 8]. However, another factor that may influence worker capabilities has received considerably less attention: the biomechanical development after eclosion [32, 34, 35, 42]. Biomechanical factors are likely relevant, because outside foraging tasks are associated with substantial mechanical demands: biting, piercing, sucking and material transportation by flight or terrestrial locomotion all require sufficiently large muscle forces and a robust skeleton to transmit these forces without inflicting damage. To investigate how muscle forces and skeletal rigidity change in the days after eclosion, we here conduct a study of the biomechanical development of a musculoskeletal apparatus that is of particular importance in foraging, in a social insect where the mechanical demands on it are particularly strong: the head capsule of leaf-cutter ants.

Leaf-cutter ant foraging involves the cutting and transport of leaf fragments from fresh vegetation surrounding the nest [43, 44, see Fig. 1]. To meet the high mechanical demands of plant cutting, leaf-cutter ants have evolved the ability to generate excessively large weight-specific bite forces [45], involving a metabolic scope close to that measured for insect flight [46]. The ability to partake in foraging thus depends in part on the physiology of the mandible closer and opener muscles and the mechanical robustness of the skeletal system. Previous work suggested that the musculoskeletal bite apparatus of ants may not have fully matured at the time of eclosion: tooth hardness of leaf-cutter ant mandibles increases nearly threefold in the days after eclosion, a development correlated with corresponding changes in zinc-concentration [34]; and the cephalic muscles of *Pheidole* ants grow substantially post eclosion [32]. Both results suggest the possibility of age-related constraints on foraging ability based on cuticle and muscle development.

**Figure 1.**
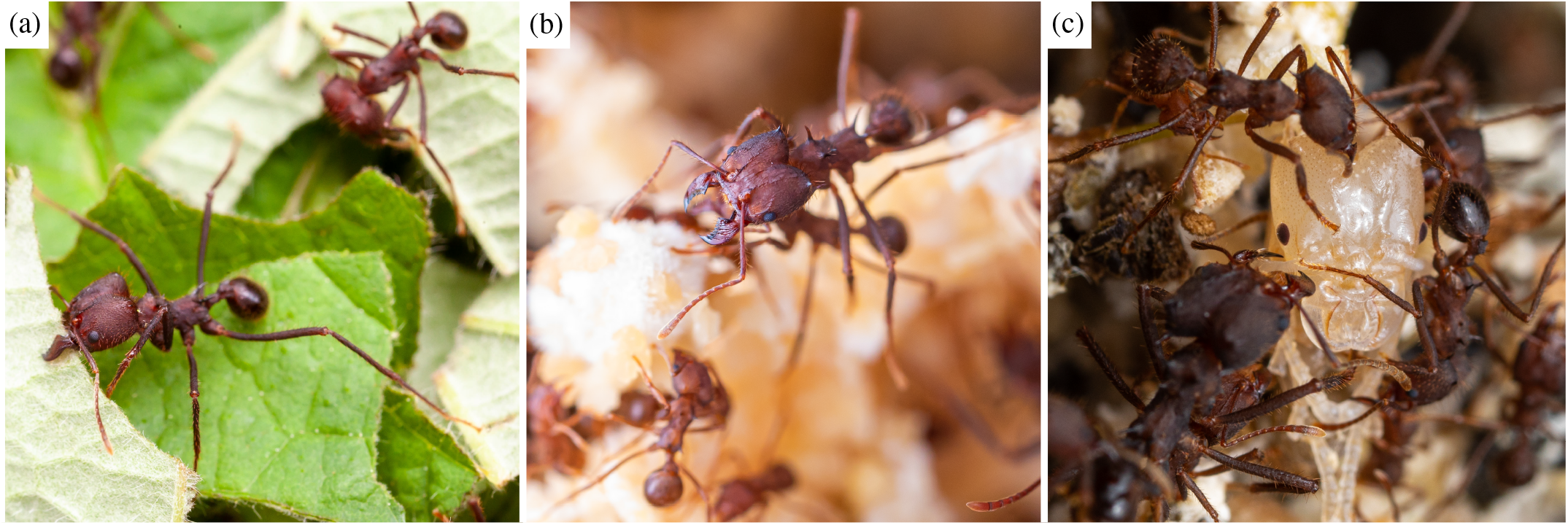
In leaf-cutter ant colonies, foraging tasks are only performed by workers exceeding a minimum age after eclosion **(a)**, whereas younger workers stay inside the nest and care for fungus **(b)** and pupae **(c)**. To investigate if this behavioural transition may be driven by a change in ability to meet the high mechanical demands of foraging, we studied the biomechanical development of the musculoskeletal bite apparatus from freshly-eclosed callows to fully-matured foragers. Photo credit: Samuel T. Fabian.

Our study builds on this work by quantifying the biomechanical development of the musculoskeletal bite apparatus of *Atta vollenweideri* leaf-cutter ants; we investigated to which extent mechanical ability varies in the days following eclosion by (i) measuring peak voluntary bite forces with a custom-built force setup; (ii) quantifying the volume of the mandible closer muscle and the thickness of the head capsule using *μ*CT imaging; and (iii), extracting the indentation modulus of the cuticle via nanoindentation experiments. By establishing a biomechanical paradigm to investigate age-related changes in the ability to partake in foraging tasks, we hope to increase our understanding of the physical constraints that may contribute to age polyethism in insects.

## Materials & methods

### Study animals

We used workers from three *Atta vollenweideri* leaf-cutter ant colonies (‘*E*’, ‘*D*’ and ‘*F*’), all founded and collected in Uruguay in 2014. The colonies were fed with bramble, cornflakes and honey water *ad libitum*, and kept under a 12 h:12 h light:dark cycle at 25°C and 50 % humidity in a climate chamber. In order to minimise confounding effects due to worker size differences [45, 47], we only collected small workers with a body mass between 3 and 7 mg, representing the lower end of forager sizes in *A. vollenweideri* [2.5-26.9 mg, see 48]. We collected 49 ants: 14 fully-darkened workers from the foraging area (n_*F*_ = 14), and 35 callows of varying cuticle brightness from the fungal gardens (n_*D*_ = 7, n_*E*_ = 8, n_*F*_ = 20). Callows were extracted by scooping fresh fungus from boxes that appeared to contain a large amount of brood into a separate container, from which those ants with visibly brighter cuticle were carefully extracted using insect tweezers. We selected ants such that there was no significant difference in mean body mass between callows (4.5*±*1.2 mg) and foragers (4.8*±*1.2 mg; Wilcoxon rank sum test: W = 214.5, p = 0.51). However, body mass may increase during maturation as a result of tissue growth (see results), which would render a size-independent comparison between callows and foragers difficult. We hence extracted a second size-metric from tomographic scans (see below), the distance between the mandibular joint centres [as defined in 47] – a metric that we considered unlikely to change with age. For callows, the joint distance was 1.13*±*0.12 mm, not significantly different to that of foragers (1.20*±*0.17 mm, Two sample t-test: t_8_ = −0.78, p = 0.46). We hence assume that any change of body mass during maturation is negligible. We combined data from all three colonies as previous work demonstrated that bite forces are consistent across *A. vollenweideri* colonies [45].

### Bite force measurements

In order to quantify bite performance, we measured the maximum bite force using a custom-built setup described in detail in Püffel et al. [45]. In brief, individual ants were held in front of two bite plates using insect tweezers (see Fig 2b). The ants then readily bit onto the two bite plates, both protruding from two mechanically uncoupled beams. The first beam can rotate about a pivot and then pushes onto a capacitive force sensor, which is thus compressed when a force is applied to the bite plate. The second beam remains stationary, such that the distance between the two outer surfaces of both bite plates – the mandibular gape required to bite – is approximately constant at 0.5 mm, or roughly a third of the average head width. Each measurement was terminated after at least five complete bite cycles or a total bite duration of more than 10 s. For the youngest and supposedly weakest callows, this condition was difficult to verify from the force trace alone, as peak bite forces were below 10 mN, equivalent to about twice the sensor noise (see results). For these ants (n = 11), we identified bites based on direct observations via a top-down camera, which filmed the ants during the experiment with 30 fps. After the bite force experiment, all ants were weighed (AX304 Microbalance, 310 g x 0.1 mg, Mettler Toledo, Greifensee, Switzerland), and sacrificed by freezing.

**Figure 2.**
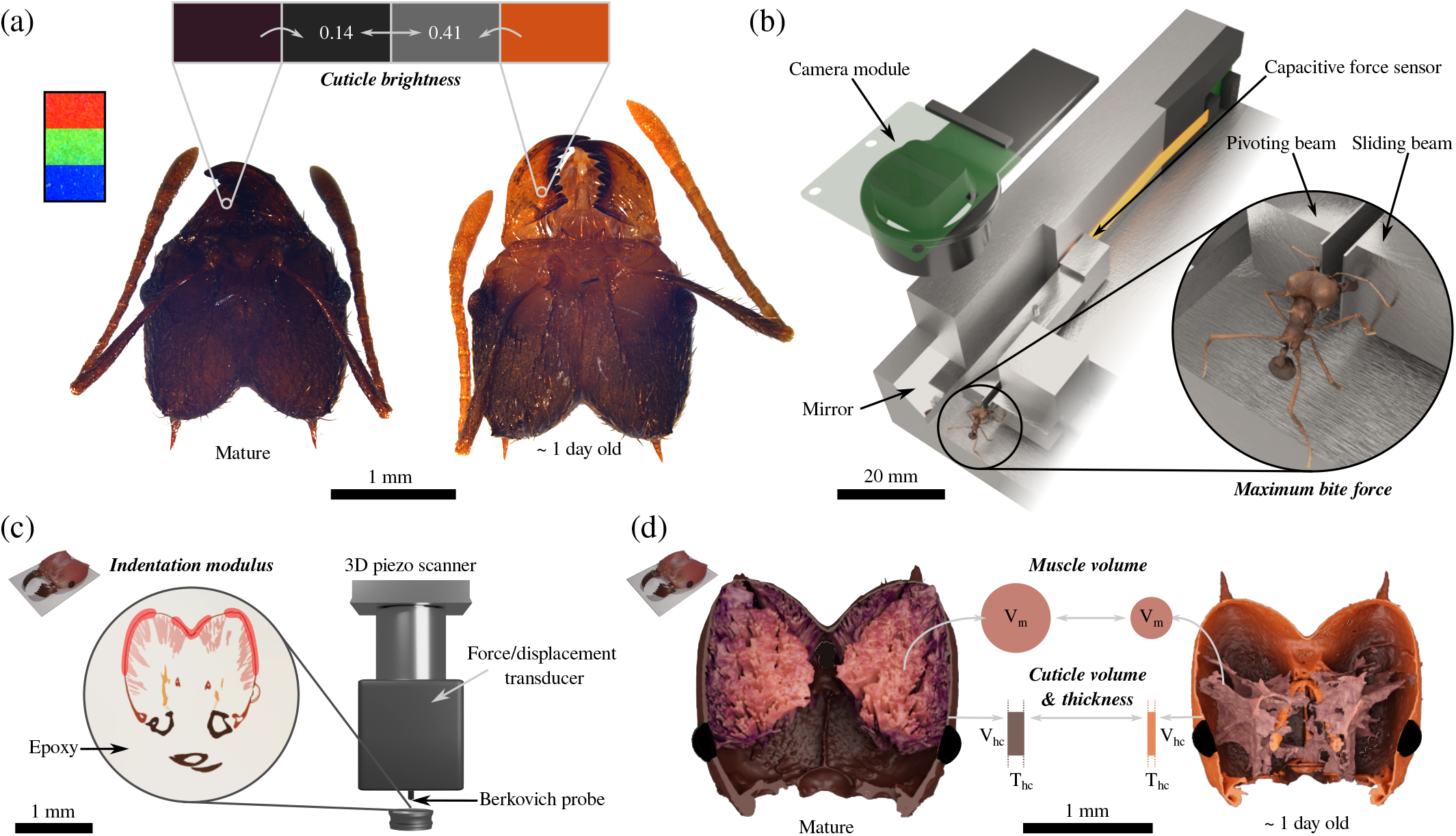
In order to study the biomechanical development of the musculoskeletal bite apparatus post eclosion, we extracted adult *Atta vol-lenweideri* leaf-cutter ants at different time points after eclosion from the foraging area and fungal gardens of three different colonies. We quantified *cuticle brightness, maximum bite force, indentation modulus, muscle volume* and *cuticle volume & thickness*. **(a)** as proxy for age post eclosion, we measured the brightness of the mandible cuticle from standardised photographs, both for randomly selected ants, and for a smaller subset of ants of known age after eclosion. **(b)** We quantified bite performance with a custom-built force sensor [described in detail in 45, 3D model of biting ant created by Fabian Plum with the open-source photogrammetry platform ‘scAnt’, see 49]. **(c - d)** In order to assess the change in structural rigidity of the head capsule, we measured its indentation modulus via nanoindentation experiments at several locations in the horizontal head plane (red areas), and its average thickness, *T*_*hc*_, from *μ*CT images. From these images, we also extracted the volumes of the mandible closer muscle and head cuticle, *V*_*m*_ and *V*_*hc*_, respectively.

From the recorded bite force traces, the maximum bite forces were extracted. However, these force maxima were still influenced by head orientation and bite location both on the mandible and bite plate. In order to remove the influence of these factors, we calculated the maximum bite force at an equivalent mandible outlever, the most distal end of the mandible, using the procedure described in detail in Püffel et al. [45]. We did not correct for differences in mandibular opening angle as this requires assumptions on muscle physiology [see 45]. However, the opening angles of callow and forager bites were almost identical, 78*±*6° versus 77*±*5° (Two sample t-test: t_47_ = 0.63, p = 0.53), so that any confounding effects are likely unsystematic. Bite forces of mature *A. vollenweideri* ants are maximal at an opening angle of about 60°; the forces measured here are about 15 % lower than this peak [45].

### Cuticle brightness

In order to approximate the age of the collected workers post eclosion, we measured cuticle brightness [e. g. 16, 22, 34], which typically decreases after eclosion, in concert with cuticle sclerotisation [50, and see below]. To make this qualitative link quantitative, we also extracted the cuticle brightness from a set of nine callows of known age post eclosion [also see 32]. To this end, late-stage pupae were taken from the fungal gardens and placed in a separate container (≈ 15 × 8 × 5 cm) with a small amount of fungus and dozens of minims to maintain them. The pupae were checked daily, and as soon as the legs had unfolded, they were marked with paint [Edding 4000 paint marker, Edding AG, Ahrensburg, Germany, see 42], and sacrificed after one (n = 3, 4.1*±*0.9 mg), three (n = 3, 4.1*±*0.8 mg) or five further days in the container (n = 3, 4.5±1.0 mg; the difference in body mass between these subsets was not significant, ANOVA: F_1,6_ = 0.22, p = 0.81).

To quantify cuticle brightness, we photographed worker head capsules in the horizontal plane using a high-resolution light microscope equipped with an apochromatic lens (camera: DMC5400, microscope: Z6 Apo controlled via *LAS X*; Leica Microsystems GmbH, Wetzlar, Germany; see Fig. 2a). Heads were placed on white paper next to a printed RGB colour stripe, which served as a baseline to enable comparison between relative colour differences across photographs (see Fig. 2a). To minimise such differences, we used the same exposure time and colour profile in *LAS X*, and kept lighting conditions approximately constant, using the microscope-internal light source and two external LED lamps. From each image, the RGB values for a defined set of ‘regions of interest’ (ROIs) were extracted using the ‘Color histogram’ function in *Fiji* [51]. A rectangular ROI was taken from each colour stripe (red, green, blue), approximately spanning 200 × 150 pixels, and one circular ROI from a ‘tooth-free’ region of the mandible blade with a diameter of 30 pixels (see Fig 2a).

The perceived colour of the mandible is likely affected by both pigmentation and variations in translucency of the cuticle due to local variations in thickness. To minimise confounding effects, we always measured cuticle brightness at locations where the left and right mandible did not overlap. We then calculated cuticle brightness from the RGB values as *b*_*RGB*_ = (*R* + *G* + *B*)*/*(3 · 255), equivalent to an unweighted greyscale conversion native to *Fiji*.

Neither body mass nor the brightness of the colour stripe differed significantly between monitored callows (n = 9) and all other workers (n = 49; body mass: Wilcoxon rank sum test: W = 197.5, p = 0.63; brightness: Wilcoxon rank sum test: W = 374, p = 0.18). However, we found a significant negative correlation between body mass and cuticle brightness across the ants used for bite experiments (Spearman’s rank correlation: *ρ*_47_ = −0.38, p < 0.01), such that dark ants (*b*_*RGB*_ *<* 0.20) weighed 5.2*±*1.2 mg, and bright ants (*b*_*RGB*_ *>* 0.35) weighed 4.2*±*1.2 mg, approximately 20 % less. This effect may be attributed to size-dependent differences in mandible thickness, which resulted in age-independent differences in cuticle translucency and brightness. We however argue that any confounding effects of body mass are small in comparison to the effects of ageing and the associated development of the bite apparatus (see below).

### Nanoindentation

To investigate if the material properties of the head capsule change in the days post eclosion, we conducted nanoindentation experiments; this technique involves pushing a small probe with well-defined geometry into a material while simultaneously recording both force and displacement. Material properties such as the indentation modulus and indentation hardness can then be extracted from the relation between force and displacement, using established mechanical theory [52]. We used a subset of twelve ants from the 49 biting ants for the experiments, selected to cover the range of observed cuticle brightness: eight callows (4.3*±*1.2 mg) and four foragers (4.3*±*1.2 mg; the difference in body mass was not significant, Two sample t-test: t_10_ = 0.03, p = 0.97). To expose a smooth cross-section of the head capsule, the samples were embedded in two-part epoxy (EPO-Set, MetPrep Ltd., Coventry, UK), so that the dorsal head plane faced upwards (see Fig. 2c), and subsequently ground and polished. To facilitate head capsule alignment, an insect pin was pierced into each head capsule prior to embedding. After curing for at least 6 h, the samples were ground (Saphir 250 A2-ECO, QATM, Mammelzen, Germany) using abrasive silicon-carbide paper of increasing grit numbers (400, 800, 1200, 2500, 4000) in single pressure mode at 25 N for 30, 90, 120, 180 and 180 s, respectively. All samples were then polished with 0.3 *μ*m alumina and 0.06 *μ*m colloidal silica suspensions, respectively (MetPrep Ltd., Coventry, UK), at 15 N for 120 s each.

Indentations were performed with a Hysitron TriboIndenter (Ti 950, Bruker Corporation, Billerica, MA, USA) and a Berkovich probe (a three-sided pyramid manufactured from diamond). Each sample was indented in a single cross-sectional plane, but numerous times (19*±*9), and in three different regions where the closer muscle attaches (see red areas in Fig. 2c). The minimum distance between indents was approximately 20 *μ*m; all indents were at least ≈ 2 μm away from the cuticle-epoxy interface. We used a trapezoidal loading profile in closed-loop displacement control, with a load time of 5 s, a peak displacement of 300 nm, a hold time of 20 s, and an unloading time of 5 s. The indentation modulus was extracted from the unloading curve using the native control software and Oliver-Pharr analysis [see 53]; the tip-area function of the Berkovich tip was calibrated with 100 indents on fused quartz, and confirmed on polycarbonate standards supplied by the manufacturer. All measurements were conducted at ambient conditions.

### Tomography and morphometric analysis

In order to quantify muscle volume and the thickness of the head capsule, an additional subset of ten ants was prepared for *μ*CT imaging: seven callows (4.2*±*1.3 mg) and three foragers (4.7*±*1.5 mg); the difference in body mass was not significant (Two sample t-test: t_8_ = −0.60, p = 0.56). For mature workers, the labrum and antennae were removed with forceps, and about five additional holes were pierced into the head capsule using insect pins to facilitate the fixative penetration. For the callows, only the antennae were removed, and the head capsule was otherwise left intact; this precaution was necessary to minimise deformation of the head capsule during manual manipulation (see discussion). All heads were fixed in paraformaldehyde solution (4 % in PBS, Thermo Fisher Scientific, Waltham, MA, USA) for 18 h, and subsequently dehydrated via storage in 70 %, 80 %, 90 % and 100 % ethanol for one hour each.

Prior to scanning, the samples were stained with 1 % iodine in ethanol for 48-168 hours [see 54], rinsed, and transferred to a pipette tip with 95 % ethanol (for more details, see SI). The samples were imaged via X-ray microscopy (XRM), using a lab-based Xradia Versa 520 (Carl Zeiss XRM Inc., Dublin, CA, USA; with a tube voltage of 70 kV, current of 85 *μ*A, and exposure time of 500 ms), a CCD detector system with scintillator-coupled visible light optics, and a tungsten transmission target. A low energy filter was placed in the beam path (LE1, proprietary Carl Zeiss microscopy filter), and a total of 2401 projections were captured at a 4x lens magnification with 2x binning over a ‘180 degrees plus fan angle’ range. The tomograms were reconstructed from 2D projections using a commercial software package (XMReconstructor, Carl Zeiss XRM Inc., Dublin, CA, USA), with a cone-beam reconstruction algorithm based on filtered back-projection, resulting in 8-bit greyscale image stacks with isotropic voxel sizes between 2.6 to 3.4 *μ*m, or about 15 % of the smallest average measured for head capsule thickness (see results).

From the tomographic scans, the mandible closer muscle and head capsule were segmented in ITKSNAP [v 3.6, 55], and the respective tissue volumes were directly exported from the software (see Fig. 2d). The average cuticle thickness was obtained via the ‘LocalThickness’ function [boneJ plugin in *Fiji*, 56], performed on the image stack of the head capsule segmentation. To quantify the error of the thickness measurement, we created an artificial 3D image of a hollow cylinder in python [v 3.9.7, 57], with a uniform shell thickness of 10 px, within the range of average values extracted in this study (5-12 px). The estimated shell thickness was virtually uniform across the cylinder, and within ≈ 1 % of its true value. However, there may be a second source of error, arising from partial volume effects of the tomographic images; partial volume effects occur at the interfaces between two materials when a voxel is partially filled by both, resulting in an intermediate greyscale value [58]. Such effects may be particularly problematic because the lowest measured cuticle thickness was only 5 px. During segmentation, we tried to avoid the ‘smearing out’ of tissue by excluding the voxels that had greyscale values closer to those of the tissue surroundings [also see 59]. However, this process is subjective and its accuracy is difficult to quantify. We hence offer a second argument in support of the accuracy of our measurements and thus of the conclusions we draw from them: We photographed the head cross-sections used for nanoindentation with the micro-scope and camera system internal to the Hysitron TriboIndenter. We then measured cuticle thickness at 30 different locations across the sample (see red areas in Fig. 2c), as the length of the shortest line connecting both tissue boundaries. The extracted cuticle thickness of dark foragers exceeded that of the brightest callows (*b*_*RGB*_ *>* 0.35) by a factor of 2.3 (14.3*±*2.9 *μ*m vs 6.2*±*1.9 *μ*m), close to the ratio obtained from the segmented tomography scans (1.9, see results). We note that the thickness measured using light microscopy was generally lower than the average value obtained via tomography, which was ‘biased’ upwards by thickened regions around the mandible articulation (see SI Fig. 1f & g).

## Statistical analysis

To test for significant correlation between cuticle brightness and the other experimental quantities, we performed Spearman’s rank correlation tests to account for non-normality of brightness values (Shapiro-Wilk normality test: W_48_ = 0.92, p < 0.01). Due to methodological limitations, data for indentation modulus and cuticle thickness were unpaired and only available for a small subset of ants. To estimate both for all ants used in bite experiments, we characterised the relationship between indentation modulus, head capsule thickness and cuticle brightness via Ordinary Least Squares (OLS) regressions on log_10_-transformed data, and then used these regressions to estimate missing values; log-transformation was necessary, as only then were all assumptions of a linear model met [see 60].

## Results and discussion

The behavioural transition from within-nest to outside-foraging tasks with age is well established in social insects [e. g. 9, 17, 22], but it remains unclear *why* it occurs. From a biomechanical perspective, foraging requires the ability to generate and withstand substantial forces, be it for object grasping, transport, or mechanical processing. However, biomechanical traits are rarely studied in the context of age polyethism [e. g. 32, 34, 35]. To test if the mechanical demands of foraging limit the ability of young workers to partake in it, we quantified a set of key performance parameters of the musculoskeletal bite apparatus of *A. vollenweideri* leaf-cutter ants. Bite forces of fully-matured foragers exceeded those of freshly-eclosed callows by more than one order of magnitude. This variation may arise because the mandible closer muscle is not yet fully developed early after eclosion, or because the mechanical stability of the head capsule limits the maximum force that can be applied to it without collapsing. In the following paragraphs, we discuss evidence for both hypotheses, and embed our findings in the context of age-related foraging in leaf-cutter ants.

### Co-development of bite force and muscle post eclosion

One of the key performance metrics of the musculoskeletal apparatus is the maximum bite force it can generate. Across workers with different cuticle brightness, maximum bite force varied by a factor of 13, from 102*±*46 mN for dark foragers (*b*_*RGB*_ = 0.17*±*0.03) to only 8*±*6 mN for the brightest callows (*b*_*RGB*_ *>* 0.35, see Fig. 3a), barely exceeding the sensor noise.

**Figure 3.**
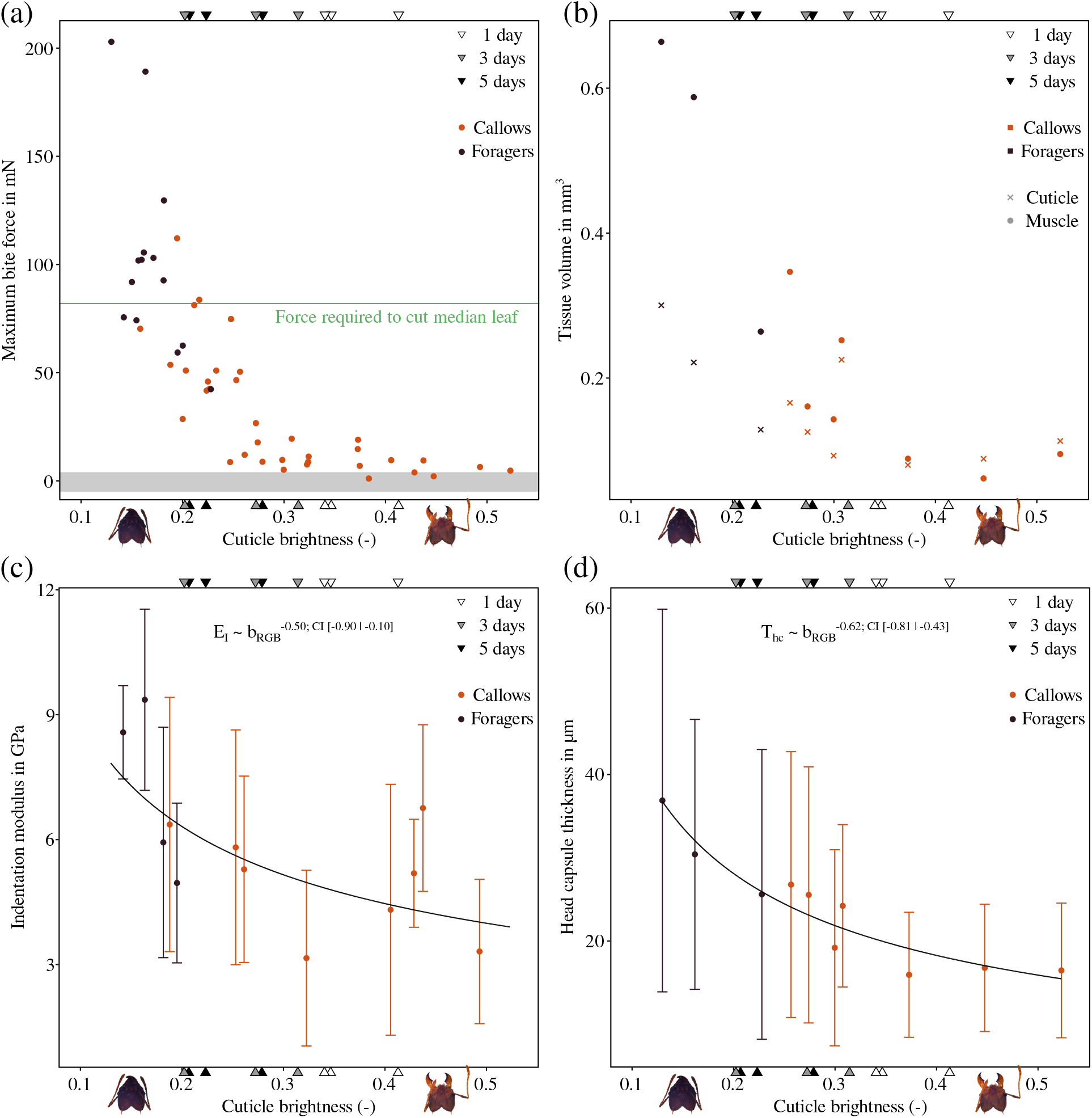
*Atta vollenweideri* leaf-cutter ant workers with varying cuticle brightness were extracted from the colonies; workers with bright or dark cuticle were extracted from the fungal gardens (callows), or from the foraging area (foragers), respectively. To link brightness to age, a set of nine late-stage pupae were photographed one, three or five days post eclosion (triangles). **(a)** Maximum bite force negatively correlates with cuticle brightness, and decreases steeply from a maximum of more than 100 mN for workers with dark cuticle to a minimum of less than 10 mN for bright callows, just in excess of sensor noise (shaded area, n = 49). The poor bite performance of callows likely constrains their ability to partake in foraging activities such as leaf-cutting; the force required to cut the median tropical leaf is about 80 mN [leaf data extracted from 61], a factor of 10 larger than the maximum bite forces callows can produce. **(b)** In order to investigate the origin of the variation in bite force, the total volume of the mandible closer muscle (circles), and of the head capsule cuticle (crosses) were extracted from segmented tomography scans. Muscle volume decreases significantly by a factor of six, and cuticle volume by a factor of two between fully-matured foragers and the brightest callows (n = 10). **(c)** To determine the flexural rigidity of the head capsule, we measured both indentation modulus of the cuticle and its thickness. The indentation modulus, *E*_*I*_, decreases significantly with cuticle brightness, *b*_*RGB*_, by a factor of 1.5 (n = 12; Ordinary Least Squares (OLS) regression on log_10_-transformed data, slope: −0.50, 95% CI: [−0.90 | −0.10], R^2^ = 0.43). **(d)** The average head capsule thickness, *T*_*hc*_, decreases from ≈ 30 μm for foragers to ≈15 *μ*m for the brightest callows (n = 10; OLS regression on log_10_-transformed data, slope: −0.62, 95% CI: [−0.81|-0.43], R^2^ = 0.87); the similar relative increase in cuticle volume and thickness suggests that most of the variation in volume stems from changes in thickness.

The cuticle brightness of these ants ranged from a minimum of 0.13 to a maximum of 0.52. For comparison, the cuticle brightness of the monitored ants with known age decreased significantly from 0.37*±*0.04 at one day to 0.24*±*0.04 at five days post eclosion (ANOVA: F_1,6_ = 6.84, p < 0.05, see Fig. 3). These results suggest that the youngest biting callows were presumably younger than 24 h, and fully-darkened foragers were older than five days, which may be a conservative estimate given that closely-related *A. sexdens rubipilosa* ants spend 3-4 weeks in the callow phase [see 34].

The change in bite force is associated with a substantial increase in the volume of the mandible closer muscle; from bright callows (*b*_*RGB*_ *>* 0.35) to foragers, muscle volume increases by a factor of six from 0.08*±*0.02 mm^3^ to 0.51*±*0.21 mm^3^ (see Table 1 and Fig. 3b; muscle volume was combined for both head hemispheres). To put this change into perspective, the muscle volume of callows is about equal to that of fully-matured workers which are four times lighter [1.1 mg, see 47, and Fig. 4a]; a size that would typically not engage in leaf-cutting [48].

**Table 1.** Results of Spearman’s rank correlation on a set of biomechanical parameters paired with cuticle brightness, *b*_*RGB*_. The degrees of freedom (Df = n-2), correlation coefficients (*ρ*) and p-values are provided for each test. All parameters apart from 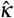 were significantly negatively correlated with cuticle brightness; 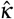 represents a proxy for cuticle strain based on maximum bite force, and the flexural rigidity and thickness of the head capsule (see main text for details).

| Parameter | Symbol | Df | $\rho$ | P-value |
| --- | --- | --- | --- | --- |
| Maximum bite force | $F_b$ | 47 | -0.89 | < 0.001 |
| Muscle volume | $V_m$ | 8 | -0.92 | < 0.001 |
| Cuticle volume | $V_{hc}$ | 8 | -0.71 | < 0.05 |
| Cuticle thickness | $T_{hc}$ | 8 | -0.94 | < 0.001 |
| Indentation modulus | $E_I$ | 10 | -0.62 | < 0.05 |
| Cuticle 'deformation' | $\kappa$ | 47 | -0.37 | < 0.01 |
| Cuticle 'strain' | $\hat{\kappa}$ | 47 | 0.02 | = 0.89 |

**Figure 4.**
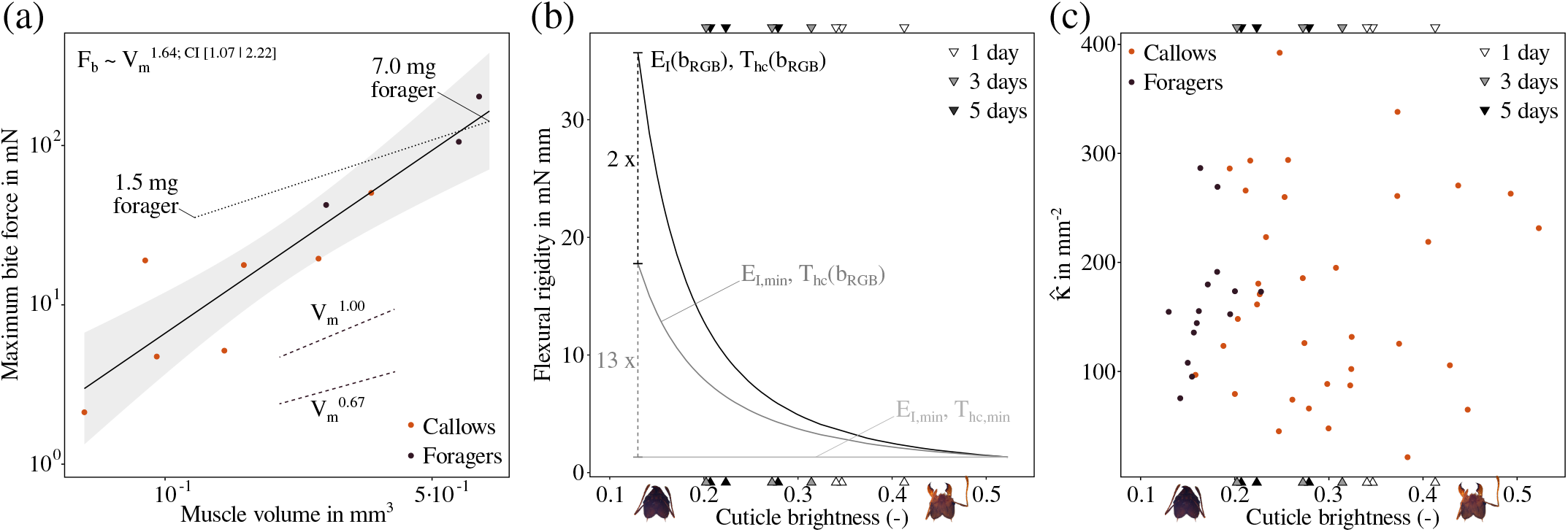
**(a)** From callows to fully-darkened foragers of the same size, the maximum bite force increased with muscle volume as *F*_*b*_ ∝ *V*_*m*_^1.64^ (OLS regression on log_10_-transformed data, 95% CI slope: [1.07 | 2.22], R^2^ = 0.85). This positive allometry suggests that both muscle volume and volume-specific bite force increase substantially post eclosion. For comparison, a fully-matured worker of 1.5 mg with the same muscle volume and third of the body weight produces four times higher bite forces [*F*_*b*_ was extracted for the same opening angle as this study, 45, 47]. **(b)** Muscle development is accompanied by changes in cuticle thickness and indentation modulus, leading to an increase of the flexural rigidity of the head capsule, *D*, by a factor 27 (black line). The majority of this increase stems from an increase in cuticle thickness, *T*_*hc*_ (factor 13.4), due the cubic dependency of *D* (see Eq. 1), whereas the indentation modulus, *E*_*I*_, contributes only linearly (factor 2.0). To visualise these relative contributions, the flexural rigidity is shown (i) as the original relationship with cuticle brightness based on the regression results for both indentation modulus and thickness (*E*_*I*_ (*b*_*RGB*_), *T*_*hc*_(*b*_*RGB*_), black line); (ii) for a variable thickness, *T*_*hc*_(*b*_*RGB*_), but constant indentation modulus, *E*_*I,min*_, fixed at its minimum (dark grey); and (iii), for constant minimum values of both in-dentation modulus and (*E*_*I,min*_, *T*_*hc,min*_, light grey). The cuticle brightness of freshly-eclosed ants of known age is shown at the top and bottom abscissa for reference (triangles). **(c)** The ratio between maximum bite force and flexural rigidity normalised with cuticle thickness, 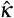, serves as a proxy for cuticle strain. 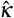 does not correlate significantly with cuticle brightness (p = 0.89, see Table 1), indicating an approximately constant mechanical demand on the head capsule during biting across the post eclosion development.

In order to assess to which extent the variation of bite force can be explained by changes in muscle volume, a theoretical prediction for the scaling relationship between both is needed. The basis for such a prediction is not obvious, because although the force of a muscle is typically proportional to its cross-sectional area [62], it is *a priori* unclear whether additional volume accumulates in area or length during muscle development. Based on geometric similarity, the cross-sectional area grows in proportion to 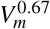, which may provide a reasonable lower bound for the expected scaling relationship. For the upper bound, we may assume that muscle fibre length remains constant, so that all volume accumulates in the crosssection, and muscle force scales in direct proportion to volume; this may be a ‘generous’ upper bound as myofibrils typically grow both in width and in length during muscle maturation [63]. Based on these assumptions, the expected scaling coefficient of bite force lies between two-thirds and one. We observed 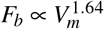, in substantial excess of both theoretical predictions (OLS regression on log_10_-transformed data, slope: 1.64, 95% CI: [1.07 | 2.22], R^2^ = 0.85; see Fig. 4a), suggesting that volume changes alone are insufficient to explain the increase of bite forces.

The mandible closer muscle in callow leaf-cutter ants is thus not only considerably smaller, but may also have a reduced stress capacity. Such an ‘underperformance’ was previously predicted by Muscedere et al. [32], who studied the development of cephalic muscles in *Pheidole* ants post eclosion; they found that freshly-eclosed callows had muscle fibres that were up to a factor of three thinner, had a non-uniform diameter, and lacked characteristic striation compared to mature ants [32]. Although we were unable to identify the ultrastructure of the mucle from the tomographic scans, we also noticed further developmental differences in addition to changes in volume: Muscle fibres of callows were less distinctly separated and often detached from the head capsule (for more details, see SI). The detachment of muscle is surprising, because muscle attachment typically matures in the early steps of muscle morphogenesis, as shown in *Drosophila* [e. g. 63, 64]. To test if muscle detachment arose as an artefact of freeze-thawing and sample preparation, we scanned another two callows (≈ 1 and 3 days old) that underwent the same protocol, but were not used in bite experiments (see SI). Muscle detachment was less severe in these ants, cautiously suggesting that (i) the bite experiment may have caused some muscle fibres to detach, and (ii) muscle formation may have only just begun at the point of eclosion; we emphasise that a more thorough study on the muscle morphogenesis in *Atta* is required to draw stronger conclusions.

Substantial muscle development after eclosion is common in insects: The cross-sectional area of beetle flight muscle [65] and locust abdominal muscle [66] increases substantially. Flight muscles in bees display a sharp increase of enzyme activity in the first days after eclosion, suggesting biochemical adjustments [38–40]. During metamorphosis in *Drosophila* flies and *Manduca* moths, the developing adult flight muscles initially lack fully-matured motor neurons required for muscle activation [e. g. 67–69]. The ‘underperformance’ of muscle observed in this study is hence likely a result of the combined effects of anatomical, physiological, biochemical and neurological deficiencies. The effect of these deficiencies is rather substantial: A mature worker of 1.5 mg has the same muscle volume as a callow, but produces four times higher bite forces at a third of its body mass [45, 47, see Fig. 4a].

### Flexural rigidity and the mechanical demands on the head capsule during biting

We have demonstrated that bite forces in callow leaf-cutter ants are strongly reduced, most likely due to incomplete muscle development. Next, we turn our attention to another biomechanical parameter that determines the ability to safely apply large bite forces: the mechanical stability of the head capsule. The ant head capsule is remarkably thin: even for mature ants, it has an average thickness comparable to that of human hair (see Fig. 3d); it is, however, locally reinforced in regions such as the tentorium, occipital suture, and around the mandibular joints (see below and SI Fig. 1f & g). The head capsule has to resist substantial size-specific bite and muscle forces. A single muscle fibre with a diameter of 25-30 *μ*m generates forces up to 0.70 mN, and ants of the considered weight have close to 1000 closer muscle fibres, resulting in a total force of ≈ 700 mN, more than ten thousand times their body weight [see 45, 47]. To estimate the structural stability of the head capsule, we introduce the flexural rigidity, *D*, the relevant metric for thin plates deformed in bending [70]:

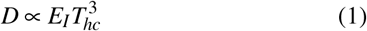

Here, *E*_*I*_ is the indentation modulus of the head capsule cuticle, and *T*_*hc*_ is its thickness.

*E*_*I*_ increased significantly with decreasing cuticle brightness (p < 0.05, see Table 1 and Fig. 3c; OLS regression slope: −0.50, 95% CI: [−0.90 | −0.10], R^2^ = 0.43). For foragers, the indentation modulus was 7.2*±*2.1 GPa, 1.5 times higher than for bright callows (*b*_*RGB*_ *>* 0.35), 4.9*±*1.5 GPa; both values are well within the range of moduli reported for insect cuticle [0.4-30 GPa, see 71, 72]. The relative increase in modulus is consistent with previous work on insects: the bending modulus of locust tibiae increases approximately threefold during the growth phase following the final moult [73]; the storage modulus of beetle elytra increases approximately sixfold in the first week after eclosion [74]; and the tooth hardness of leaf-cutter ant mandibles increases by a factor of nearly three [34], a development associated with increased zinc-concentration and possibly resistance to mandibular wear [also see 42]. In some leaf-cutter ant species, a biomineral layer forms on the epicuticle a week after eclosion, resulting an a twofold increase in hardness [75]. The biomechanical development of insect cuticle is often linked to tanning and sclerotisation, a process associated with water loss, cross-linking of cuticle proteins with chitin, and a resulting increase in modulus [50, 71, 74, 76]. Indeed, cuticle hydration affects both indentation modulus and hardness [71, 77], and because we conducted experiments of small samples in ambient conditions, we likely overestimate the modulus, and underestimate its increase during post-eclosion development [see Fig. 3 in 72, and 74].

The head capsule thickness, *T*_*hc*_, in turn increased significantly by a factor of two from 16.4*±*0.4 *μ*m for bright callows to 31.0*±*5.7 *μ*m for foragers (p < 0.001, see Table 1 and Fig. 3d; OLS regression slope: −0.62, 95% CI: [−0.81|-0.43], R^2^ = 0.87); this variation is comparable to the cuticle thickness range measured from the pronotum of other Myrmicine workers of similar head width [≈ 1.7 mm, see Fig. 2 in 78]. Notably, the increase in cuticle thickness appears to be non-uniform, indicated by the increase of the standard deviation with decreasing cuticle brightness (see Fig. 3d): For bright callows (*b*_*RGB*_ *>* 0.35), the standard deviation is 7.8*±*0.3 *μ*m, significantly smaller than for foragers, 18.9*±*3.6 *μ*m (Spearman’s rank correlation: *ρ*_8_ =-0.94, p < 0.001; this significant difference also holds for the coefficient of variation: *ρ*_8_ = −0.70, p < 0.05). The increased variability of the head capsule thickness suggests a ‘targeted’ growth of cuticle during post-eclosion development, perhaps in regions most prone to deformation [also see 79]. Cuticle volume increases significantly, too, in almost direct proportion to cuticle thickness (p < 0.05, see Table 1 and Fig. 3b): Foragers have an average cuticle volume of 0.22*±*0.09 mm^3^, exceeding that of bright callows by a factor of 2.3 (*b*_*RGB*_ *>* 0.35, 0.09*±*0.02 mm^3^). Cuticle growth post eclosion has been reported for other insects such as e. g. locusts [73, 80], grasshoppers [81], and moths [82], and is typically associated with the internal deposition of additional layers of endocuticle [73, 81]. In the initial growth phase following the final moult, locusts deposit about 1.8*μ*m of endocuticle per day [73]. Assuming that the age difference between foragers and bright callows is one week, the cuticle growth rate for leaf-cutter ants is about the same, 14.6 *μ*m / 7 d ≈ 2 *μ*m/d.

The combined effects of the changes in indentation modulus and cuticle thickness result in a drastic increase of the flexural rigidity with cuticle brightness [see Fig. 4b; to obtain a numerical value for *D* from Eq. 1, we used a proportionality constant 1*/*(12(1−0.3^2^)), see 70]. From the lightest to the darkest workers, the flexural rigidity increased by a staggering factor of 27, from 1.3 to 35.7 mN mm; we note that this result is based on interpolation via regression, and is hence affected by the associated uncertainties of the slope (see CIs in Fig. 3). The majority of this increase is driven by the change in cuticle thickness (factor of 13.4), which is cubed in Eq. 1, whereas the indentation modulus contributes linearly and thus has a much smaller effect (factor of 2.0). In order to make the implications of this difference in flexural rigidity tangible, we calculated the expected deflection for two plates of the same area, but of different thickness and moduli, approximated by *T*_*hc*_ and *E*_*I*_, respectively, for a forager with a cuticle brightness of *b*_*RGB*_ = 0.15 and a callow with *b*_*RGB*_ = 0.5. Under the same load, equal to half of the maximum muscle stress extracted for closely-related *A. cephalotes* majors [83], the resulting deflection of callow cuticle is 17 times higher than for dark cuticle; absolute values of deflection are estimated in the SI.

These results invite another hypothesis why callows ‘underperform’ when biting, in addition to continued muscle growth and physiological development (see Fig. 4a). Callows may choose to bite with sub-maximal muscle force in order to avoid large deformations of the head capsule. We consider two possible constraints on deformation: (i) absolute deformation, relevant if the elastic deformation is sufficiently large to cause damage inside the head, e. g. by compressing soft tissues; and (ii), relative deformation, relevant if the stress in the cuticle exceeds its elastic limit, causing fissures or permanent deformation. First, we approximate the absolute cuticle deformation resulting from biting, *κ*, as ratio between maximum bite force and flexural rigidity. For foragers, *κ* = 5.09*±*1.74 mm^−1^, 1.5 times higher than for bright callows *κ* = 3.36*±*1.89 mm^−1^. This dif-ference, although significant (p < 0.01, see Table 1), is remarkably small in comparison to the large variation in both bite force and flexural rigidity. Second, we estimate relative cuticle deformation or ‘strain’ as 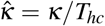. Strikingly, 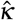 is not significantly correlated with cuticle brightness (p = 0.89, see Table 1 and Fig. 4c). suggesting that maintaining equal cuticle strain may be a constraint on maximum muscle activation. We stress that the numerical values of *κ* and 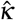 do not translate to actual cuticle deformation and strain, respectively, but are approximately proportional to them; the counter-intuitive units arise from the square of a missing length scale that causes the bending moment in the cuticle, which is likely proportional to the constant external head dimensions (also see Eq. 1 in SI).

Bite forces, muscle ultrastructure and volume, as well as head capsule rigidity all develop in the days following eclosion. Indeed, the increase of all extracted parameters with decreasing cuticle brightness follow a similar pattern, with a rapid increase across a narrow range of brightness values (see Fig. 3). This mechanical co-development results in an approximately constant mechanical strain on the head capsule cuticle associated with maximum bite force throughout the callow phase of the workers. Future work will need to address to which extent these changes are causally linked, as is observed in vertebrates [e. g. 84–86].

### Biomechanical limitations of foraging ability in leaf-cutter ants

We set out to investigate if the behavioural transition from within-nest to outside-foraging tasks may arise from variation in biomechanical performance. We have reported evidence for strong changes in bite force in the days following eclosion. Next, we address briefly if these changes may explain why young callows do not forage. In leaf-cutter ants, foraging involves the cutting and carrying of leaf fragments from fresh vegetation surrounding the nest [43]. The ability of a forager to cut leaves depends on the ratio of two key forces: the maximum bite force, and the minimum force required to initiate and propagate a cut through leaf lamina and veins [87]. Based on published mechanical data for around 1000 tropical leaves, the expected cutting forces range between 7-828 mN, with a median of 82 mN [45, 61]. Thus, the foragers used in our study would be able to cut approximately half of the measured tropical leaves, whereas bright callows could cut virtually none of them. Foraging also involves the carrying of leaf fragments: A forager of 5 mg may carry fragments of around 15 mg, or three times its body weight [48, 88]. This load may sound heavy, but the gravitational force that needs to be countered is only 0.15 mN, small even compared to the poor bite performance of callows. However, fragments cut from grasses may be as long as 30 mm [48], five times longer than the ants themselves [89], or 20 times their head length [47]; carrying fragments of this length poses a difficult mechanical challenge, which requires continuous adjustments of the neck angle to maintain stability during walking, and likely involves large moments around the neck joint [89, 90]. Assuming that neck muscles are similarly underdeveloped early after eclosion, the ability to carry large fragments may be limited in young callows; we however note that different muscle may follow different developmental agendas as shown for *Pheidole* ants [mandible closer vs antennal muscle, 32].

Our results suggest that the foraging ability of young workers is largely constrained by a poor bite performance based on underdeveloped muscles and a low mechanical stability of the head capsule – a finding that likely extends to in-nest cutting as well [see 91]. In addition, the role of mechanical constraints in influencing age polyethism may continue beyond the time frame considered here: For example, older foragers with worn mandibles may require more than twice the force to cut the same material than those with pristine mandibles [35, and Püffel et al., unpublished data], and are indeed more likely to carry leaves instead of cutting [35]; eventually, the oldest workers often switch to tasks related to waste disposal [23, 92].

## Conclusion and outlook

In the days following eclosion, the musculoskeletal bite apparatus of young leaf-cutter ants undergoes substantial biomechanical development: the maximum bite force, muscle volume, head capsule thickness and the cuticle-mechanical properties all increase substantially, from a point where bite forces are too low to cut leaves and the head capsule is too mechanically unstable to resist large muscle forces, to full biomechanical ability. These results suggest that *Atta* leaf-cutter ant callows are not yet able to partake in foraging activities involving cutting – adding direct experimental support to the hypothesis that age polyethism is largely dictated by developmental factors [e. g. 22, 24]. The extent to which callows readily perform in-nest tasks such as fungal garden or brood care, in turn, remains poorly understood [17, 18]. Future work on the behavioural development of callows inside the nest may be able to integrate the findings of this study to gain a more comprehensive understanding of the behavioural repertoire expansion of leaf-cutter ants. Our findings further suggest exciting avenues for future research on the codependency of muscle and cuticle development, which hints at an important role of mechanical stimuli, an area of research that has received considerable interest in vertebrates [e. g. 84–86], but less so in invertebrates [e. g. 63, 93]. Many tasks in colonies of social insects impose mechanical demands, and we thus hope that a biomechanical paradigm will help to increase our general understanding of age polyethism more broadly.

## Acknowledgments

We thank Samuel T. Fabian and Fabian Plum for providing the photographs and the CAD model, respectively, used in Fig. 1 and 2. This study is part of a project that has received funding from the European Research Council (ERC) under the European Union’s Horizon 2020 research and innovation programme (Grant agreement No. 851705) awarded to DL.

## References

[1] Wilson E. 1990 Success and dominance in ecosystems: the case of the social insects. Ecology Institute.

[2] Schultheiss P, Nooten SS, Wang R, Wong MK, Brassard F, Guenard B. 2022 The abundance, biomass, and distribution of ants on earth. Proceedings of the National Academy of Sciences 119: e2201550119.

[3] Oster GF, Wilson EO. 1978 Caste and ecology in the social insects. Princeton University Press.

[4] Cole BJ. 2020 Comparative advantage and caste evolution. Evolution 74: 655–659.

[5] Wilson EO. 1980 Caste and division of labor in leaf-cutter ants (Hymenoptera: Formicidae: Atta): II. the ergonomic optimization of leaf cutting. Behavioral Ecology and Sociobiology 7: 157–165.

[6] Tofts C. 1993 Algorithms for task allocation in ants. (a study of temporal polyethism: theory). Bulletin of Mathematical Biology 55: 891–918.

[7] Robinson GE, Page Jr RE, Huang Z. 1994 Temporal polyethism in social insects is a developmental process. Animal Behaviour 48: 467–469.

[8] Beshers SN, Fewell JH. 2001 Models of division of labor in social insects. Annual review of entomology 46: 413–440.

[9] Seeley TD. 1982 Adaptive significance of the age polyethism schedule in honeybee colonies. Behavioral Ecology and Sociobiology 11: 287–293.

[10] Robinson GE, Page Jr RE, Strambi C, Strambi A. 1989 Hormonal and genetic control of behavioral integration in honey bee colonies. Science 246: 109–112.

[11] Cameron SA. 1989 Temporal patterns of division of labor among workers in the primitively eusocial bumble bee, Bombus griseocollis (hymenoptera: Apidae) 1. Ethology 80: 137–151.

[12] Calderone NW. 1995 Temporal division of labor in the honey bee, Apis mellifera a developmental process or the result of environmental influences? Canadian Journal of Zoology 73: 1410– 1416.

[13] Huang ZY, Robinson GE. 1996 Regulation of honey bee division of labor by colony age demography. Behavioral Ecology and Sociobiology 39: 147–158.

[14] O’Donnell S, Jeanne RL. 1993 Methoprene accelerates age polyethism in workers of a social wasp (Polybia occidentalis). Physiological Entomology 18: 189–194.

[15] Shorter JR, Tibbetts EA. 2009 The effect of juvenile hormone on temporal polyethism in the paper wasp Polistes dominulus. Insectes Sociaux 56: 7–13.

[16] Wilson EO. 1976 Behavioral discretization and the number of castes in an ant species. Behavioral Ecology and Sociobiology 1: 141–154.

[17] Wilson EO. 1980 Caste and division of labor in leaf-cutter ants (Hymenoptera: Formicidae: Atta): I. the overall pattern in A. Sextens. Behavioral Ecology and Sociobiology 7: 143–156.

[18] Fowler HG. 1983 Alloethism in a leaf-cutting ant: laboratory studies on Atta texana (hymenoptera: Formicidae: Attini). Zoologische Jahrbücher. Abteilung für allgemeine Zoologie und Physiologie der Tiere 87: 529–538.

[19] Beshers SN, Traniello JF. 1996 Polyethism and the adaptiveness of worker size variation in the attine ant Trachymyrmex septentrionalis. Journal of Insect Behavior 9: 61–83.

[20] Julian GE, Fewell JH. 2004 Genetic variation and task specialization in the desert leaf-cutter ant, Acromyrmex versicolor. Animal Behaviour 68: 1–8.

[21] Thomas ML, Elgar MA. 2003 Colony size affects division of labour in the ponerine ant Rhytidoponera metallica. Naturwissenschaften 90: 88–92.

[22] Seid MA, Traniello JF. 2006 Age-related repertoire expansion and division of labor in Pheidole dentata (hymenoptera: Formicidae): a new perspective on temporal polyethism and behavioral plasticity in ants. Behavioral Ecology and Sociobiology 60: 631– 644.

[23] Camargo R, Forti LC, Lopes J, Andrade A, Ottati A. 2007 Age polyethism in the leaf-cutting ant Acromyrmex subterraneus brunneus forel, 1911 (hym., formicidae). Journal of Applied Entomology 131: 139–145.

[24] Muscedere ML, Willey TA, Traniello JF. 2009 Age and task efficiency in the ant Pheidole dentata: young minor workers are not specialist nurses. Animal Behaviour 77: 911–918.

[25] Vieira SA, Fernandes WD, Antonialli-Junior WF. 2010 Temporal polyethism, life expectancy, and entropy of workers of the ant Ectatomma vizottoi almeida, 1987 (formicidae: Ectatomminae). Acta ethologica 13: 23–31.

[26] Hinze B, Leuthold R. 1999 Age related polyethism and activity rhythms in the nest of the termite Macrotermes bellicosus (isoptera, termitidae). Insectes Sociaux 46: 392–397.

[27] Jeanson R. 2019 Within-individual behavioural variability and division of labour in social insects. Journal of Experimental Biology 222: jeb190868.

[28] Tofilski A. 2002 Influence of age polyethism on longevity of workers in social insects. Behavioral Ecology and Sociobiology 51: 234–237.

[29] Yanagihara S, Suehiro W, Mitaka Y, Matsuura K. 2018 Age-based soldier polyethism: old termite soldiers take more risks than young soldiers. Biology Letters 14: 20180025.

[30] Lucas C, Hughson BN, Sokolowski MB. 2010 Job switching in ants: role of a kinase. Communicative & Integrative Biology 3: 6–8.

[31] Ingram KK, Gordon DM, Friedman DA, Greene M, Kahler J, Peteru S. 2016 Context-dependent expression of the foraging gene in field colonies of ants: the interacting roles of age, environment and task. Proceedings of the Royal Society B: Biological Sciences 283: 20160841.

[32] Muscedere ML, Traniello JF, Gronenberg W. 2011 Coming of age in an ant colony: cephalic muscle maturation accompanies behavioral development in Pheidole dentata. Naturwissenschaften 98: 783.

[33] Muscedere ML, Traniello JF. 2012 Division of labor in the hyperdiverse ant genus Pheidole is associated with distinct subcaste- and age-related patterns of worker brain organization. PLoS One 7: e31618.

[34] Schofield RM, Nesson MH, Richardson KA. 2002 Tooth hardness increases with zinc-content in mandibles of young adult leafcutter ants. Naturwissenschaften 89: 579–583.

[35] Schofield RM, Emmett KD, Niedbala JC, Nesson M. 2011 Leafcutter ants with worn mandibles cut half as fast, spend twice the energy, and tend to carry instead of cut. Behavioral Ecology and Sociobiology 65: 969–982.

[36] Ben-Shahar Y, Robichon A, Sokolowski M, Robinson G. 2002 Influence of gene action across different time scales on behavior. Science 296: 741–44.

[37] Habenstein J, Thamm M, Rössler W. 2021 Neuropeptides as potential modulators of behavioral transitions in the ant Cataglyphis nodus. Journal of Comparative Neurology 529: 3155–3170.

[38] Roberts SP, Elekonich MM. 2005 Muscle biochemistry and the ontogeny of flight capacity during behavioral development in the honey bee, Apis mellifera. Journal of Experimental Biology 208: 4193–4198.

[39] Schippers MP, Dukas R, McClelland GB. 2010 Lifetime-and caste-specific changes in flight metabolic rate and muscle biochemistry of honeybees, Apis mellifera. Journal of Comparative Physiology B 180: 45–55.

[40] Skandalis DA, Roy C, Darveau CA. 2011 Behavioural, morphological, and metabolic maturation of newly emerged adult workers of the bumblebee, Bombus impatiens. Journal of Insect Physiology 57: 704–711.

[41] Robinson GE. 1998 From society to genes with the honey bee: A combination of environmental, genetic, hormonal and neurobiological factors determine a bee’s progression through a series of life stages. American Scientist 86: 456–462.

[42] Schofield RM, Nesson MH, Richardson KA, Wyeth P. 2003 Zinc is incorporated into cuticular “tools” after ecdysis: The time course of the zinc distribution in “tools” and whole bodies of an ant and a scorpion. Journal of Insect Physiology 49: 31–44.

[43] Wirth R, Herz H, Ryel RJ, Beyschlag W, Hölldobler B. 2003 Herbivory of Leaf-Cutting Ants: A Case Study on Atta colombica in the Tropical Rainforest of Panama. Berlin, Heidelberg, New York: Springer Science & Business Media.

[44] Hölldobler B, Wilson EO. 2010 The leafcutter ants: civilization by instinct. W. W. Norton & Company, New York City, NY, USA.

[45] Püffel F, Roces F, Labonte D. 2022 Strong positive allometry of bite force in leaf-cutter ants increases the range of cuttable plant tissues. bioRxiv.

[46] Roces F, Lighton JR. 1995 Larger bites of leaf-cutting ants. Nature 373: 392.

[47] Püffel F, Pouget A, Liu X, Zuber M, van de Kamp T, Roces F, Labonte D. 2021 Morphological determinants of bite force capacity in insects: a biomechanical analysis of polymorphic leafcutter ants. Journal of the Royal Society Interface 18: 20210424.

[48] Röschard J, Roces F. 2003 Fragment-size determination and size-matching in the grass-cutting ant Atta vollenweideri depend on the distance from the nest. Journal of Tropical Ecology 19: 647–653.

[49] Plum F, Labonte D. 2021 scant—an open-source platform for the creation of 3d models of arthropods (and other small objects). PeerJ 9: e11155.

[50] Politi Y, Bar-On B, Fabritius HO. 2019 Mechanics of arthropod cuticle-versatility by structural and compositional variation. In: Architectured Materials in Nature and Engineering, Springer. pp. 287–327.

[51] Schindelin J, Arganda-Carreras I, Frise E, Kaynig V, Longair M, Pietzsch T, Preibisch S, Rueden C, Saalfeld S, Schmid B, Tinevez J, White D, Hartenstein V, Eliceiri K, Tomancak P, Cardona A. 2012 Fiji: An open-source platform for biological-image analysis. Nature methods 9: 676–682.

[52] Ebenstein DM, Pruitt LA. 2006 Nanoindentation of biological materials. Nano today 1: 26–33.

[53] Oliver WC, Pharr GM. 1992 An improved technique for determining hardness and elastic modulus using load and displacement sensing indentation experiments. Journal of materials research 7: 1564–1583.

[54] Metscher BD. 2009 Microct for comparative morphology: simple staining methods allow high-contrast 3d imaging of diverse nonmineralized animal tissues. BMC physiology 9: 1–14.

[55] Yushkevich PA, Piven J, Hazlett HC, Smith RG, Ho S, Gee JC, Gerig G. 2006 User-guided 3d active contour segmentation of anatomical structures: significantly improved efficiency and reliability. NeuroImage 31: 1116–1128.

[56] Domander R, Felder AA, Doube M. 2021 Bonej2 - refactoring established research software. Wellcome Open Research 6.

[57] Van Rossum G, Drake FL. 2009 Python 3 Reference Manual. Scotts Valley, CA: CreateSpace.

[58] Withers PJ, Bouman C, Carmignato S, Cnudde V, Grimaldi D, Hagen CK, Maire E, Manley M, Du Plessis A, Stock SR. 2021 X-ray computed tomography. Nature Reviews Methods Primers 1: 1–21.

[59] Souza A, Udupa JK, Saha PK. 2005 Volume rendering in the presence of partial volume effects. IEEE Transactions on Medical Imaging 24: 223–235.

[60] Peña EA, Slate EH. 2006 Global validation of linear model assumptions. Journal of the American Statistical Association 101: 341–354.

[61] Onoda Y, Westoby M, Adler PB, Choong AMF, Clissold FJ, Cornelissen JHC, Díaz S, Dominy NJ, Elgart A, Enrico L, Fine PVA, Howard JJ, Jalili A, Kitajima K, Kurokawa H, McArthur C, Lucas PW, Markesteijn L, Pérez-Harguindeguy N, Poorter L, Richards L, Santiago LS, Sosinski Jr EE, Van Bael SA, Warton DI, Wright IJ, Joseph Wright S, Yamashita N. 2011 Global patterns of leaf mechanical properties. Ecology Letters 14: 301–312.

[62] Josephson RK. 1975 Extensive and intensive factors determining the performance of striated muscle. Journal of Experimental Zoology 194: 135–153.

[63] Lemke SB, Schnorrer F. 2017 Mechanical forces during muscle development. Mechanisms of Development 144: 92–101.

[64] Luis NM, Schnorrer F. 2021 Mechanobiology of muscle and myofibril morphogenesis. Cells & Development 168: 203760.

[65] De Kort C. 1990 Thirty-five years of diapause research with the colorado potato beetle. Entomologia Experimentalis et Applicata 56: 1–13.

[66] Rose U, Ferber M, Hustert R. 2001 Maturation of muscle properties and its hormonal control in an adult insect. Journal of Experimental Biology 204: 3531–3545.

[67] Fernandes JJ, Keshishian H. 1998 Nerve-muscle interactions during flight muscle development in Drosophila. Development 125: 1769–1779.

[68] Hughes SM, Salinas PC. 1999 Control of muscle fibre and motoneuron diversification. Current Opinion in Neurobiology 9: 54–64.

[69] Bayline RJ, Duch C, Levine RB. 2001 Nerve-muscle interactions regulate motor terminal growth and myoblast distribution during muscle development. Developmental Biology 231: 348–363.

[70] Young WC, Budynas RG. 2002 Roark’s formulas for stress and strain. McGraw-Hill, 7 edition.

[71] Vincent JF, Wegst UG. 2004 Design and mechanical properties of insect cuticle. Arthropod structure & development 33: 187–199.

[72] Labonte D, Lenz AK, Oyen ML. 2017 On the relationship between indentation hardness and modulus, and the damage resistance of biological materials. Acta Biomaterialia 57: 373–383.

[73] Parle E, Taylor D. 2017 The effect of aging on the mechanical behaviour of cuticle in the locust Schistocerca gregaria. Journal of the Mechanical Behavior of Biomedical Materials 68: 247–251.

[74] Scalet JM, Sprouse PA, Schroeder JD, Dittmer N, Kramer KJ, Kanost MR, Gehrke SH. 2022 Temporal changes in the physical and mechanical properties of beetle elytra during maturation. Acta Biomaterialia 151: 457–467.

[75] Li H, Sun CY, Fang Y, Carlson CM, Xu H, Ješovnik A, Sosa-Calvo J, Zarnowski R, Bechtel HA, Fournelle JH, et al. 2020 Biomineral armor in leaf-cutter ants. Nature Communications 11: 1–11.

[76] Wappner P, Quesada-Allué LA. 1996 Water loss during cuticle sclerotization in the medfly Ceratitis capitata is independent of catecholamines. Journal of insect physiology 42: 705–709.

[77] Klocke D, Schmitz H. 2011 Water as a major modulator of the mechanical properties of insect cuticle. Acta biomaterialia 7: 2935–2942.

[78] Peeters C, Molet M, Lin CC, Billen J. 2017 Evolution of cheaper workers in ants: a comparative study of exoskeleton thickness. Biological Journal of the Linnean Society 121: 556–563.

[79] Blanke A, Watson PJ, Holbrey R, Fagan MJ. 2017 Computational biomechanics changes our view on insect head evolution. Proceedings of the Royal Society B: Biological Sciences 284: 20162412.

[80] Hepburn H, Joffe I. 1974 Locust solid cuticle - a time sequence of mechanical properties. Journal of Insect Physiology 20: 497–506.

[81] Neville A. 1963 Daily growth layers for determining the age of grasshopper populations. Oikos : 1–8.

[82] Hopkins TL, Kramer KJ. 1992 Insect cuticle sclerotization. Annual Review of Entomology 37: 273–302.

[83] Püffel F, Johnston R, Labonte D. 2023 A biomechanical model for the relation between bite force and mandibular opening angle in arthropods. Royal Society Open Science 10: 221066.

[84] Nowlan NC, Murphy P, Prendergast PJ. 2008 A dynamic pattern of mechanical stimulation promotes ossification in avian embryonic long bones. Journal of Biomechanics 41: 249–258.

[85] Nowlan NC, Dumas G, Tajbakhsh S, Prendergast PJ, Murphy P. 2012 Biophysical stimuli induced by passive movements compensate for lack of skeletal muscle during embryonic skeletogenesis. Biomechanics and Modeling in Mechanobiology 11: 207– 219.

[86] Boselli F, Freund JB, Vermot J. 2015 Blood flow mechanics in cardiovascular development. Cellular and Molecular Life Sciences 72: 2545–2559.

[87] Clissold F. 2007 The biomechanics of chewing and plant fracture: Mechanisms and implications. Advances in Insect Physiology 34.

[88] Rudolph SG, Loudon C. 1986 Load size selection by foraging leaf-cutter ants (Atta cephalotes). Ecological Entomology 11: 401–410.

[89] Moll K, Roces F, Federle W. 2010 Foraging grass-cutting ants (Atta vollenweideri) maintain stability by balancing their loads with controlled head movements. Journal of Comparative Physiology A 196: 471–480.

[90] Moll K, Roces F, Federle W. 2013 How load-carrying ants avoid falling over: mechanical stability during foraging in Atta vol- lenweideri grass-cutting ants. Public Library of Science One 8: e52816.

[91] Garrett RW, Carlson KA, Goggans MS, Nesson MH, Shepard CA, Schofield RM. 2016 Leaf processing behaviour in Atta leafcutter ants: 90% of leaf cutting takes place inside the nest, and ants select pieces that require less cutting. Royal Society Open Science 3: 1–12.

[92] Hart AG, Ratnieks FL. 2001 Task partitioning, division of labour and nest compartmentalisation collectively isolate hazardous waste in the leafcutting ant Atta cephalotes. Behavioral Ecology and Sociobiology 49: 387–392.

[93] Parle E, Dirks JH, Taylor D. 2017 Damage, repair and regeneration in insect cuticle: The story so far, and possibilities for the future. Arthropod Structure & Development 46: 49–55.

