## Supplementary Information for "Developmental biomechanics and age polyethism in leaf-cutter ants"

### Supplementary Materials

#### Further notable differences between callows and foragers

In addition to the quantitative differences detailed in the main text, we noticed a number of qualitative differences between callows and foragers: The mandibles of callow ants sometimes remained partially open after sacrificing them by freezing, whereas foragers always had their mandibles maximally closed; this difference presumably reflects differences in muscle development. During dissection, callow head capsules were noticeably ‘squishy’, and considerable amounts of watery liquid were expelled upon deformation (also see methods). When examined under the microscope, the mandible closer muscle of matured foragers appeared to fill the majority of the head capsule, whereas callows had much smaller muscles; consistent with $\mu$CT data. Other differences may include the presence of a large amount of adipocytes surrounding callow head muscle, as observed for *Pheidole* and *Monomorium* ants [1, 2]; however, we were unable to derive robust conclusions on the existence of such fat bodies in the muscle tissue of callow *Atta* using the available CT images.

#### Plate model

We estimate the deformation of the head capsule caused by muscle contraction using a simple mechanical model. We model the head capsule as a plate of area, $A = H_w/4 \cdot H_h/2$, where head width, $H_w$ and head height, $H_h$, were extracted for a 4.2 mg *Atta vollenweideri* ant [3, see Fig. 1]. The thickness of the plate is set equal to the average local thickness of the head capsule, $T_{hc}$, and the modulus is approximated as $E_I$ (both measured in this study, see results). For simplicity, we assume a quadratic shape of the plate with a side length of $a = \sqrt{A}$, clamped along all sides. In fully-matured leaf-cutter ants, the closer muscle attaches to the internal side of the head capsule with a maximum theoretical density of 50 % [assuming a cylindrical shape, see 3]. At full muscle activation, the area load on the plate is thus equal to half of the maximum muscle stress, $\sigma = \sigma_m/2$ [$\sigma_m$ = 1.16 MPa, extracted for *A. cephalotes* majors in 4]. From plate theory, the associated maximum deflection then follows as:

$$w = \alpha\sigma \frac{a^4}{E_I T_{hc}^3} \tag{1}$$

where the geometry factor is $\alpha = 0.0138$ for square plates [5]. For a forager with a cuticle brightness of 0.15, $E_I = 7.3$ GPa and $T_{hc} = 34\,\mu$m, so that the maximum deflection is 1.4 $\mu$m or 4 % of the cuticle thickness. For a callow with a cuticle brightness of 0.50, $E_I = 4.0$ GPa and $T_{hc} = 16\,\mu$m, the deflection is 24.0 $\mu$m, equivalent to 150 % of the cuticle thickness. We hence argue that the deflection of the head capsule during muscle contraction is likely within the elastic limit for matured foragers, whereas it may cause critical damage for callows exposed to the same hypothetical muscle force. Assuming a much smaller area load of $\sigma = \sigma_m/26$, equivalent to the observed difference in bite force between foragers and bright callows, the deflection is only 1.8 $\mu$m for callows, or 12 % of the cuticle thickness.

### References


[1] Jensen P, Børgesen L. 2000 Regional and functional differentiation in the fat body of pharaoh’s ant queens, *Monomorium pharaonis* (l.). *Arthropod Structure & Development* **29**: 171–184.

[2] Muscedere ML, Traniello JF, Gronenberg W. 2011 Coming of age in an ant colony: cephalic muscle maturation accompanies behavioral development in *Pheidole dentata*. *Naturwissenschaften* **98**: 783.

[3] Püffel F, Pouget A, Liu X, Zuber M, van de Kamp T, Roces F, Labonte D. 2021 Morphological determinants of bite force capacity in insects: a biomechanical analysis of polymorphic leaf-cutter ants. *Journal of the Royal Society Interface* **18**: 20210424.

[4] Püffel F, Johnston R, Labonte D. 2023 A biomechanical model for the relation between bite force and mandibular opening angle in arthropods. *Royal Society Open Science* **10**: 221066.

[5] Young WC, Budynas RG. 2002 Roark’s formulas for stress and strain. McGraw-Hill, 7 edition.

[6] Domander R, Felder AA, Doube M. 2021 Bonej2 - refactoring established research software. *Wellcome Open Research* **6**.

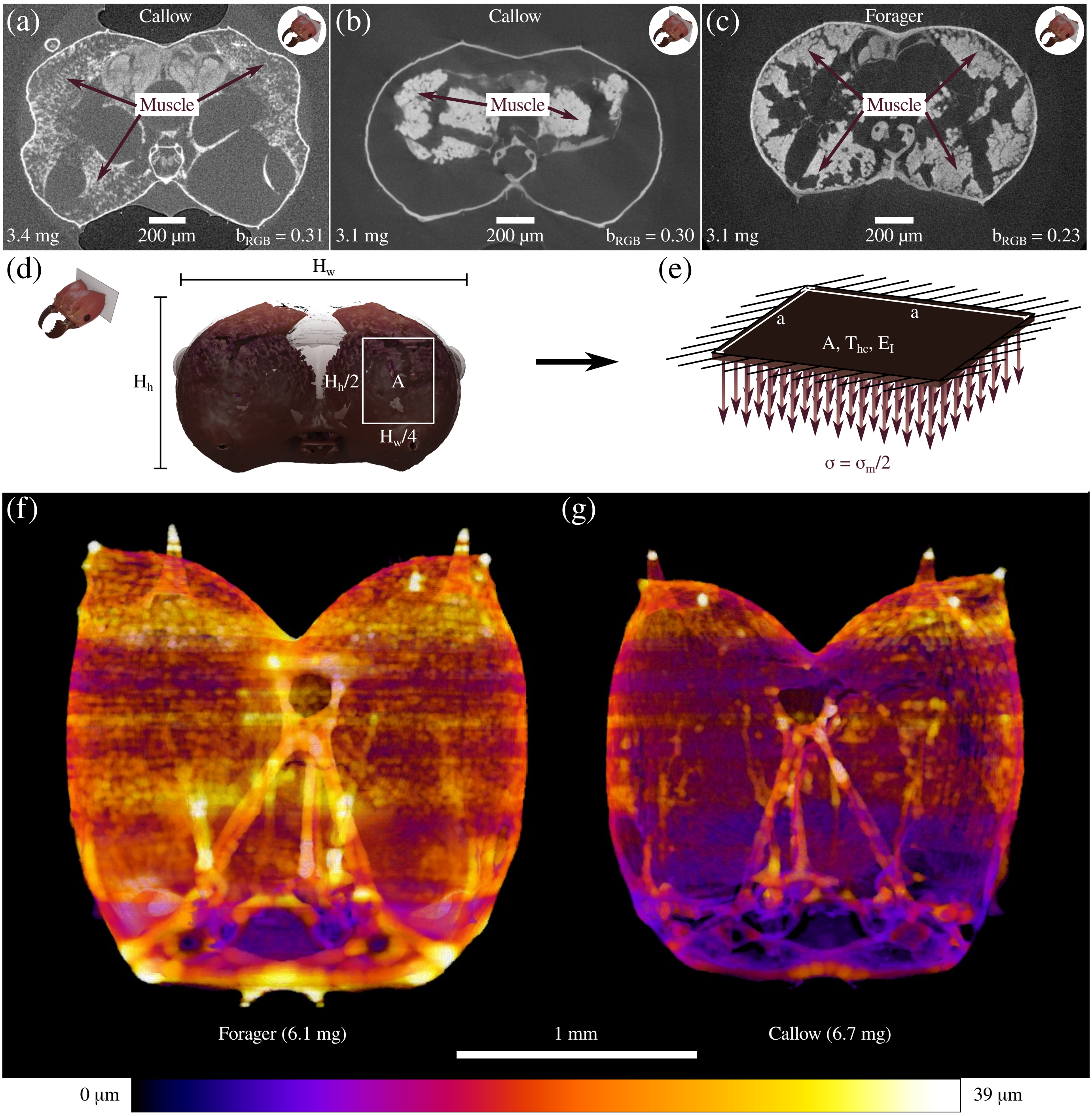


Figure 1 | **(a-c)** Prior to $\mu$CT imaging, *Atta vollenweideri* leaf-cutter ants of varying cuticle brightness, $b_{RGB}$, were used for bite experiments, sacrificed by freezing, photographed and prepared for scanning. **(b)** We observed that the closer muscle tissue of callows that were used in bite experiments was often detached from the inner surface of the head capsule. **(a)** To test if this observation was an artefact of repeated freeze-thawing, we scanned another two callows that underwent the same treatment before tomography, but were not used in bite experiments; muscle fibres of these ants showed less detachment from the head capsule. **(c)** In fully-matured foragers, muscle detachment was not observed. **(d-e)** In order to illustrate the effects of cuticle maturation on head capsule deformation, we modelled the head capsule as a square plate with an area, $A = H_w/4 \cdot H_h/2$, where head width and height, $H_w$ and $H_h$, respectively, were extracted for a 4.2 mg *A. vollenweideri* ant [see 3], a side length $a = \sqrt{A}$, a thickness, $T_{hc}$, and a modulus, $E_I$. The plate was clamped at all sides, and a uniform area load, equal to half of the maximum muscle stress, $\sigma_m$, extracted for closely-related *A. cephalotes* majors [4], was applied. For the same load case, the resulting maximum deflection for a bright callow would be 24.0 $\mu$m or 1.5 times the cuticle thickness, 17 times higher than for a mature forager. In other words, if callows generated bite forces as large as those of fully-matured foragers, their head capsule would deform considerably, potentially resulting in permanent damage. **(f-g)** The thickness of the head capsule cuticle was measured via the 'LocalThickness' function [boneJ plugin in *Fiji*, 6]. The local thickness varies considerably across the head capsule, where brighter values correspond to thicker regions. Both average thickness and the coefficient of variation were significantly higher in foragers **(f)** compared to callows **(g)**, suggesting a 'targeted' deposition of endocuticle during post-eclosion development (for more details, see main text).
